## Supplemental figures for "Focused ultrasound excites neurons via mechanosensitive calcium accumulation and ion channel amplification"

### Supplementary Figures, Movie and Tables

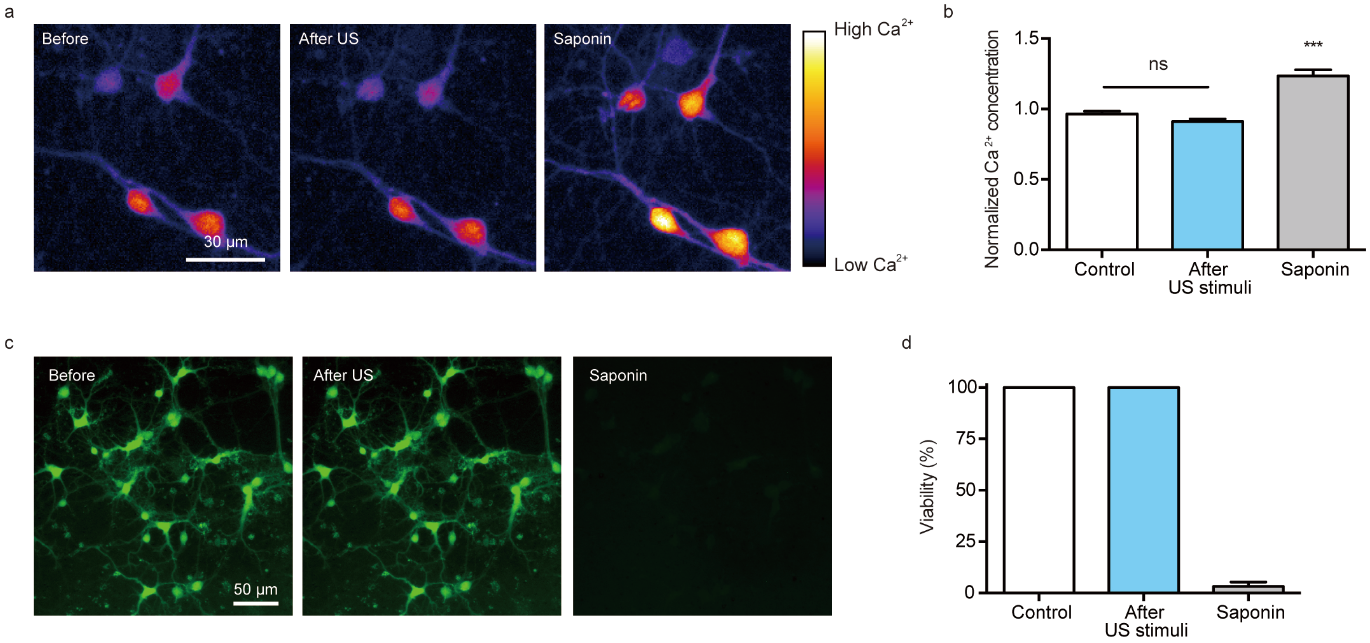


**Supplementary Figure 1 | Safety test at cellular level after ultrasound stimulation.** (a) Change of GCaMP6f baseline intensity before and after ultrasound stimuli (15 W/cm$^2$, 30 times every 20 sec), and (b) their quantification (n=117 from 2 dish, one way ANOVA followed by Tukey's multiple comparison test, p=0.0587 (control vs after US)). (c) Live neuron images before and after US stimulation and saponin treatment (after 5 mins from 100 µg/ml saponin treatment). And (d) quantification of neuron viability before and after the ultrasound stimuli and saponin treatment (n=10 ROI from 2 dishes).

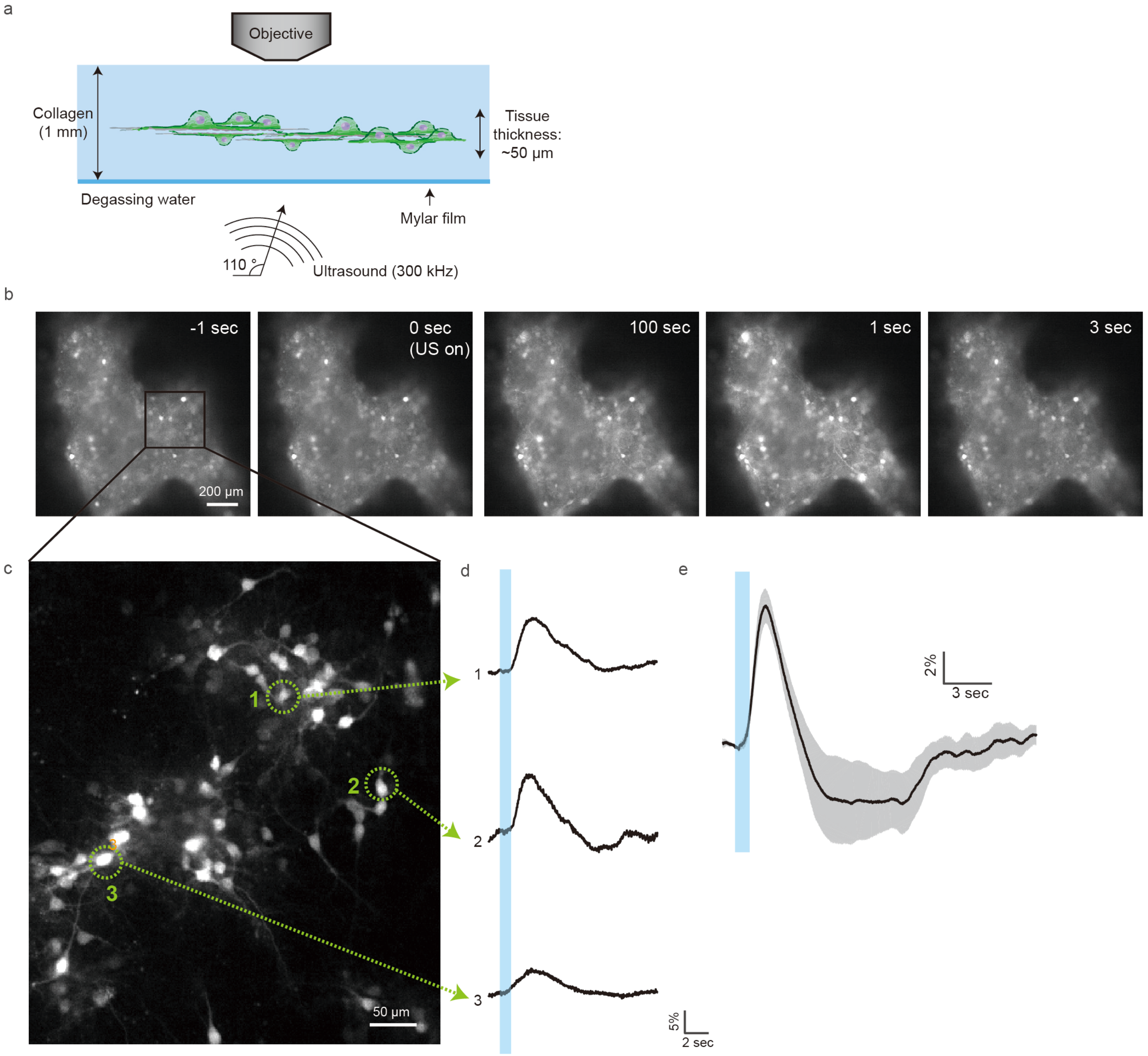


**Supplementary Figure 2 | Ultrasound stimulation to 3D neural tissue model embedded in collage hydrogel.** (a) Schematics of ultrasound stimulation to 3D collagen tissue model. (b) Time lapse images of calcium responses to ultrasound stimulation from the 3D cultured GCaMP6f neurons. (c) A snapshot of the 3D cultured GCaMP6f neurons and (d) Calcium responses from selected neurons. (e) Averaged calcium response (n= 4 dishes). Mean trace in solid and SEM is shaded.

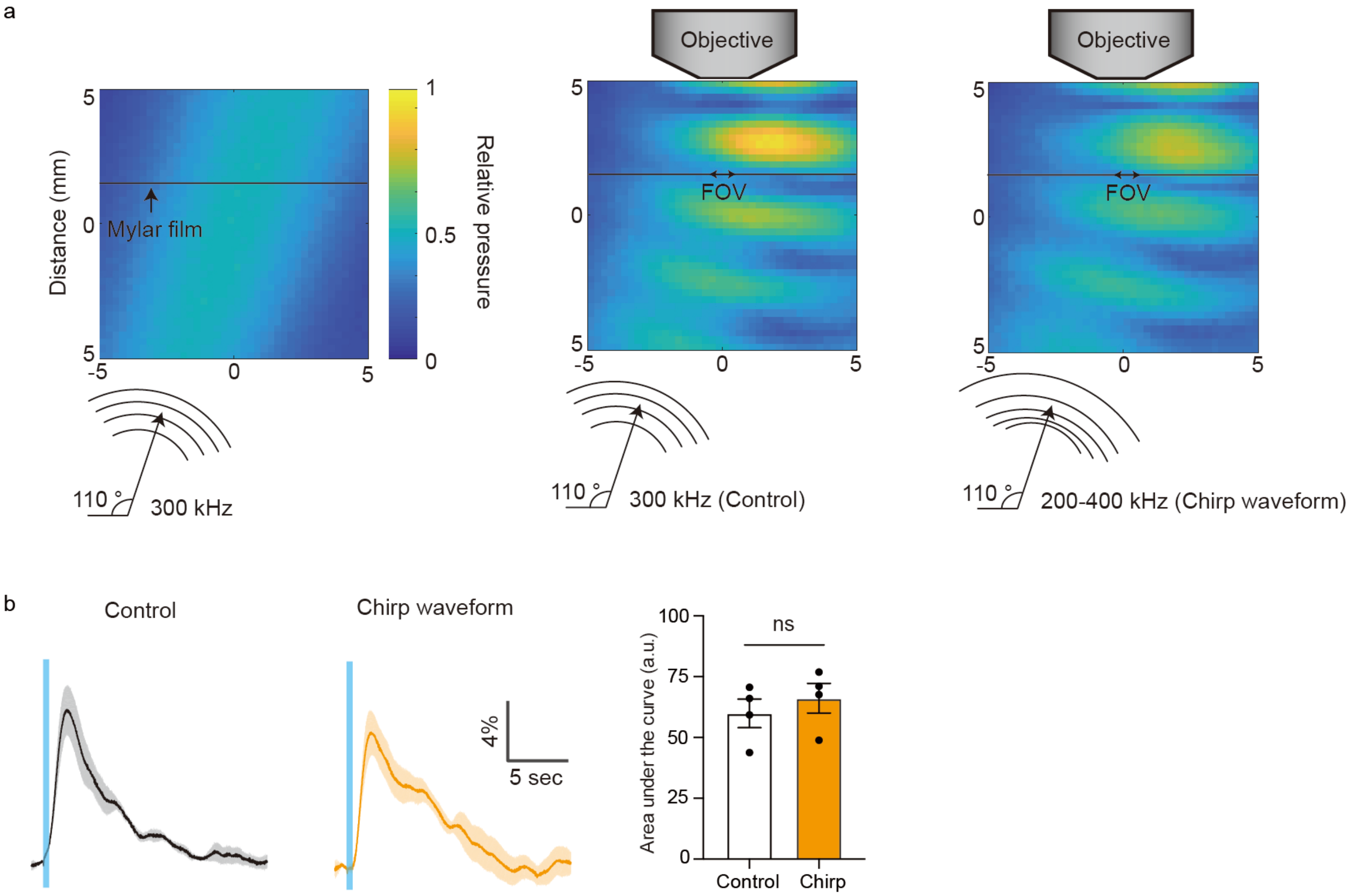


**Supplementary Figure 3 | Ultrasonic neuron stimulation with chirp waveform.** (a) Pressure profiles measured using a fiber optic hydrophone in three configurations. Left: deep underwater with no acoustic reflectors (i.e. "free field") with 300 kHz ultrasound. Middle: air/water interface and microscope objective lens at same locations as used in fluorescent imaging studies with 300 kHz ultrasound. Right: same configuration as middle with chirp waveform (200-400 kHz ultrasound, 10 µs sweep time). Standing wave pattern produced by reflection off objective lens decreased with chirp waveform compared to control waveform. (b) Calcium responses to ultrasound with normal and chirp waveform and their (c) quantifications (n= 4 dishes, Paired T-test, p=0.4891). Mean trace in solid and SEM is shaded.

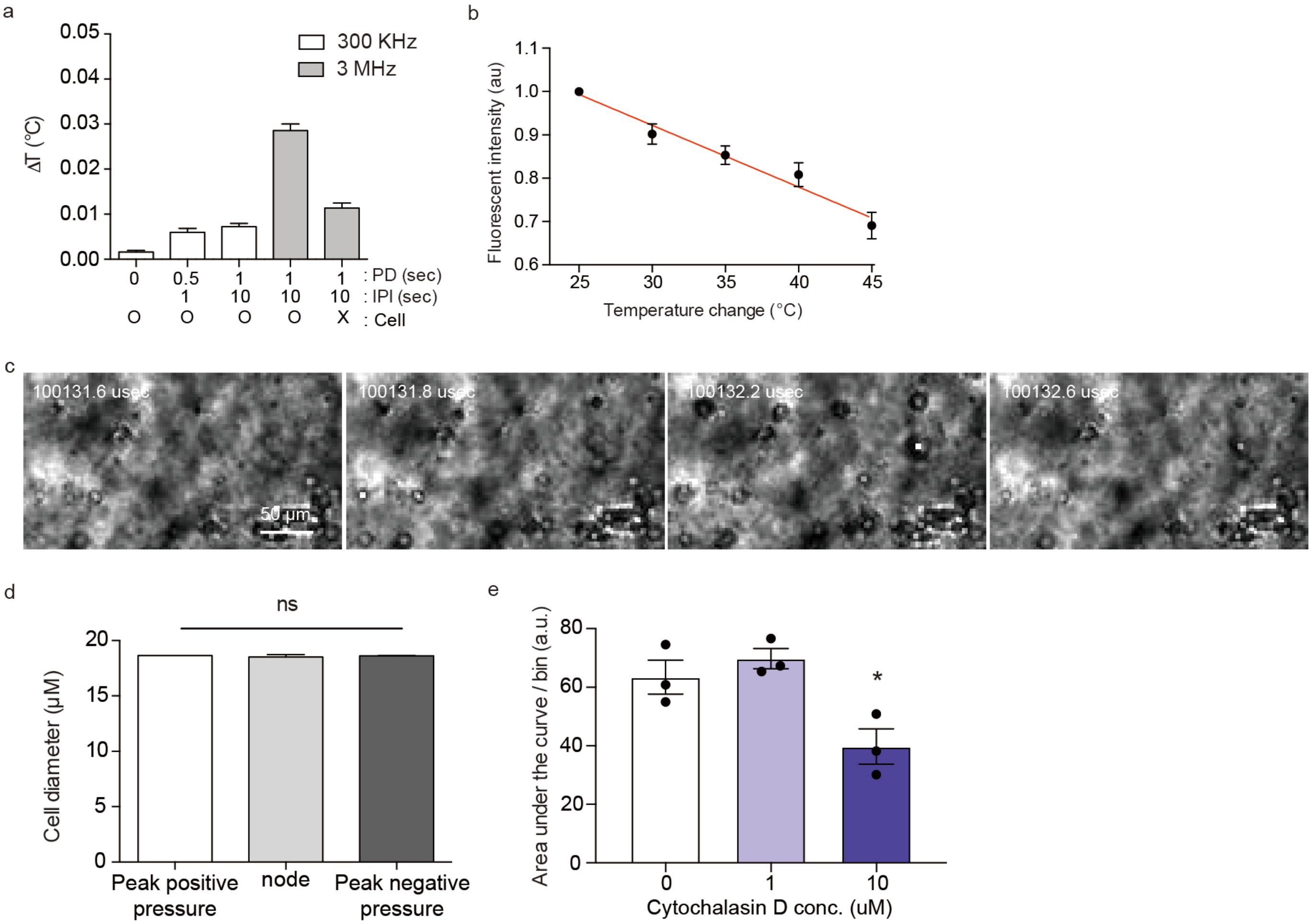


**Supplementary Figure 4 |** (a) Temperature measurement using an optic hydrophone near neurons during ultrasound stimulation with various parameters (n=20, 15 and 40 W/cm$^2$ intensities were used for 300 kHz and 3 MHz, respectively). (b) Fluorescence intensity of mCherry decreased as increasing the temperature (n=20, 2%/°C, R square=0.9240). (c) Ultra-high-speed imaging of microbubbles (bubble size=4 μm) while ultrasound stimulation. (d) Quantification of cell diameter change during ultrasound stimulation (n=15, Paired T-test, p=0.3343). (e) Quantification of spontaneous activity before and after the actin depolymerization (bin =5 sec, n=3 dishes each, p=0.6859 (0 vs 1 μM)) with different concentrations of depolymerizers.

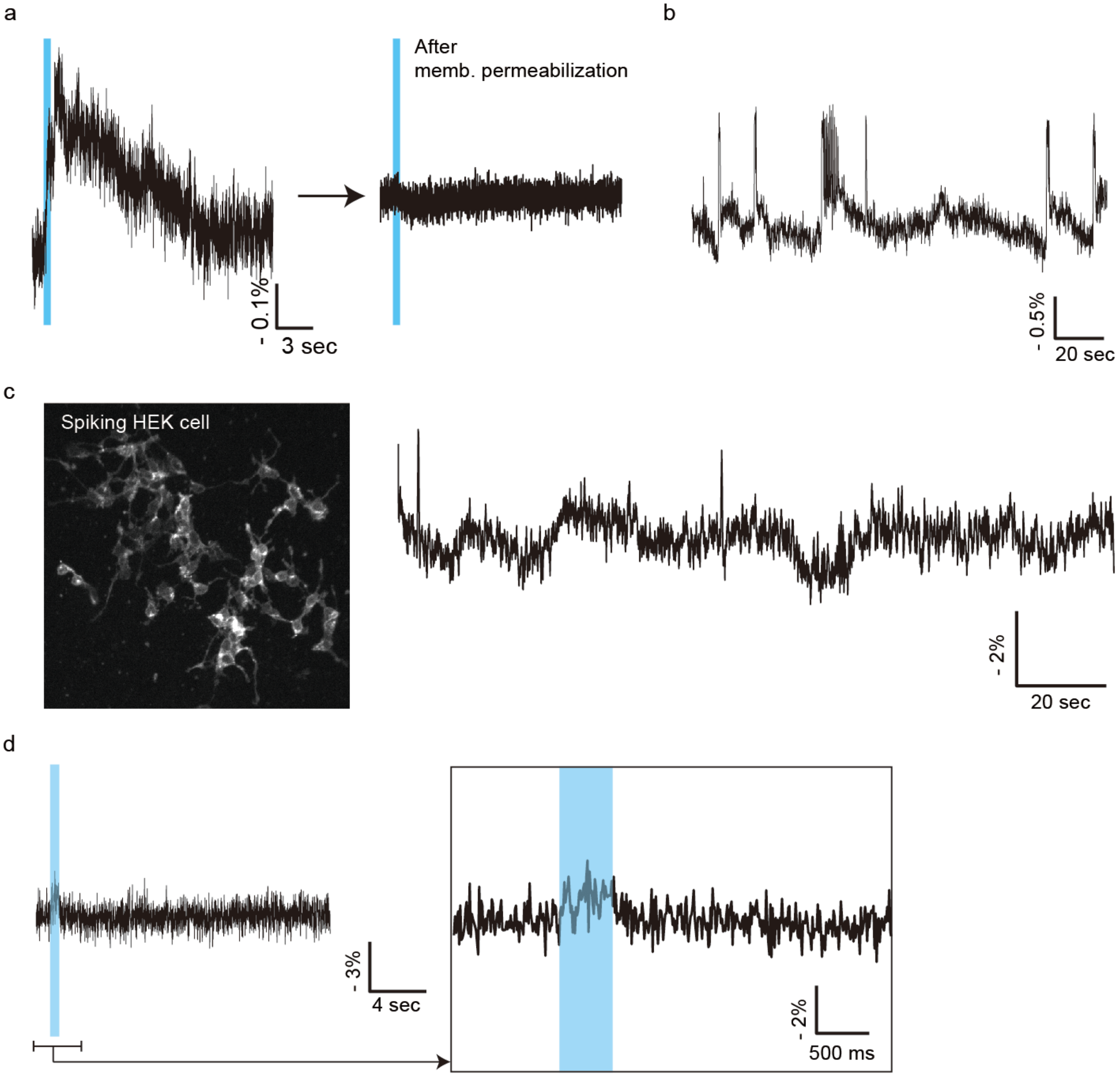


**Supplementary Figure 5 |** (a) Voltage responses to ultrasound (n= 4 dishes), and the responses disappeared after membrane permeabilization by saponin. (b) Spontaneous activity of Ace2N-expressing neurons in the absence of extracellular calcium. 1 μM bicuculine (GABAa blocker) was added to induce the hyper excitation (n=101 from a dish). (c) Ace2N-expressing spiking HEK cells and their spontaneous activity n=52 from a dish. (d) Voltage imaging of the spiking HEK cells during ultrasound stimulation, n= 82 cells from 2 dishes.

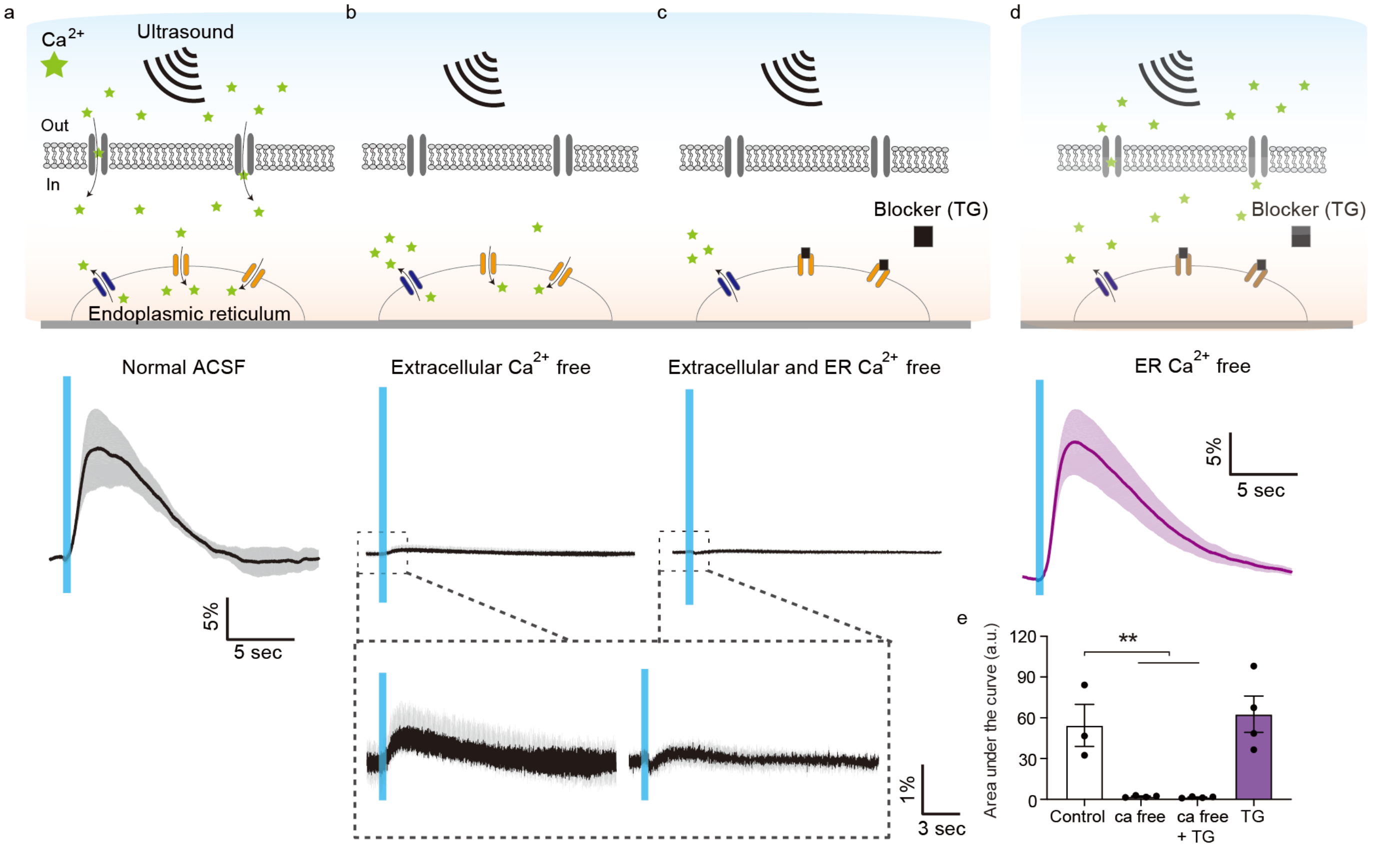


**Supplementary Figure 6 | Ultrasound stimulation triggers calcium release from endosomal reticulum.** (a) Calcium response to ultrasound stimulation in the presence of extra- and intercellular calcium (normal ACSF) (b) Calcium response to ultrasound stimulation in the absence of extracellular calcium (calcium free ACSF). (c) Calcium response to ultrasound stimulation in the absence of calcium in extracellular space and endoplasmic reticulum (calcium free ACSF + Thapsigargin). (d) Calcium response to ultrasound stimulation in the absence of calcium in endoplasmic reticulum (normal ACSF + Thapsigargin). (e) Quantification of area under the curve from each condition (n= 3 dishes for control, = 4 each for others, Unpaired T test,). Mean trace in solid and SEM is shaded.

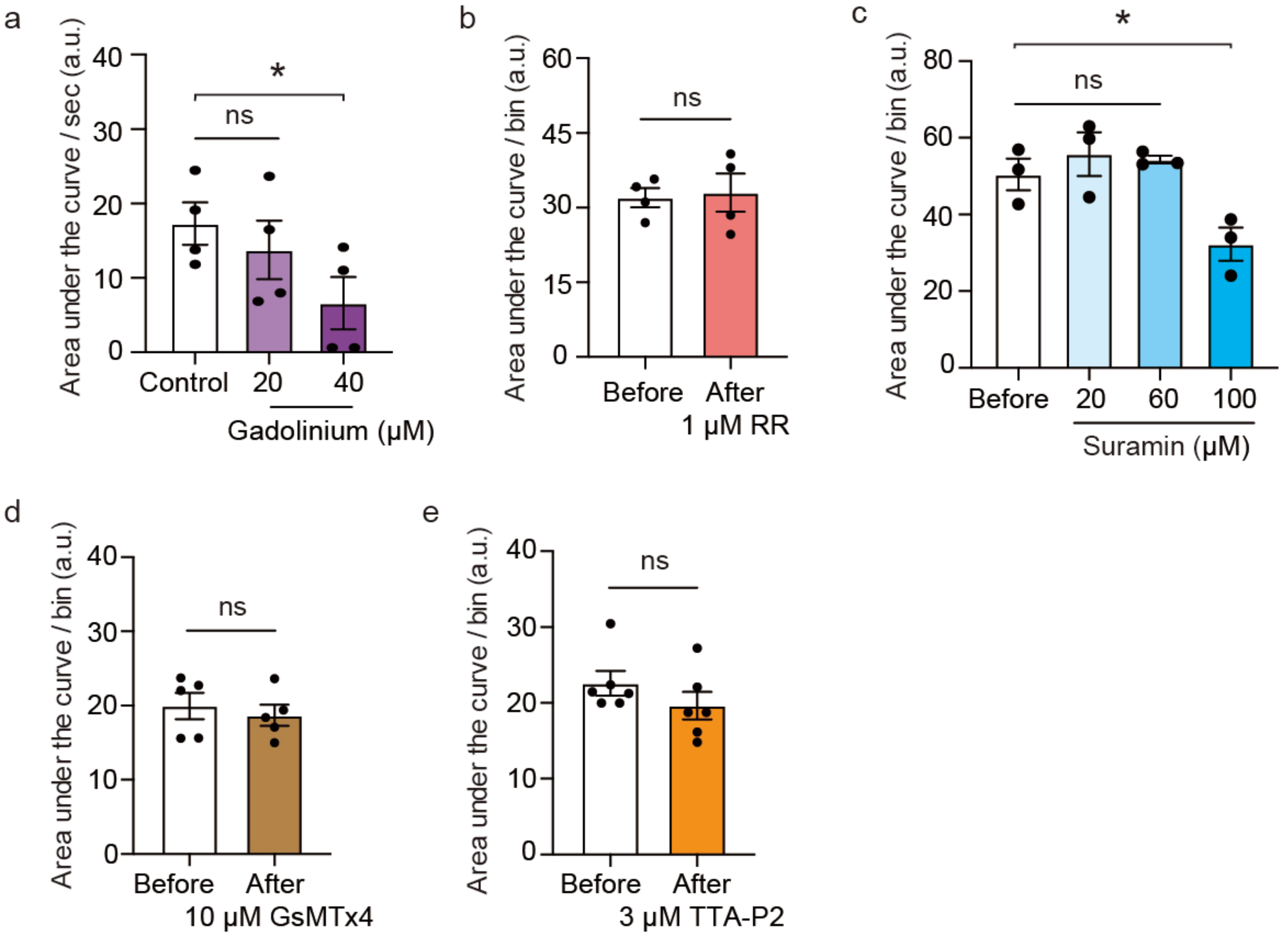


**Supplementary Figure 7 | Spontaneous activity changes before and after channel blocking.** (a) Spontaneous activity change after gadolinium treatment to block the global mechxnosensitive channels (n= 4 dishes, Paired T-test, p=0.6258 (control vs 20 uM)). (b) Spontaneous activity change after ruthenium red treatment to block the TRPV1, 2 and 4 channels (n= 4 dishes, Paired T-test, p=0.8337 . (c) Spontaneous activity change after suramin treatment to inhibit the GPCRs (n=3 dishes each, Unpaired T-test, p=0.4167 (control vs 60 uM)). (d) Spontaneous activity change after GsMTx4 treatment to inhibit the Piezo1 and TRPC1 channels (n= 5 dishes, Unpaired T-test, p=0.6005). (e) Spontaneous activity change after TTA-P2 treatment to blocker the t-type calcium channels (n= 6 dishes, Paired t test, p=0.2546).

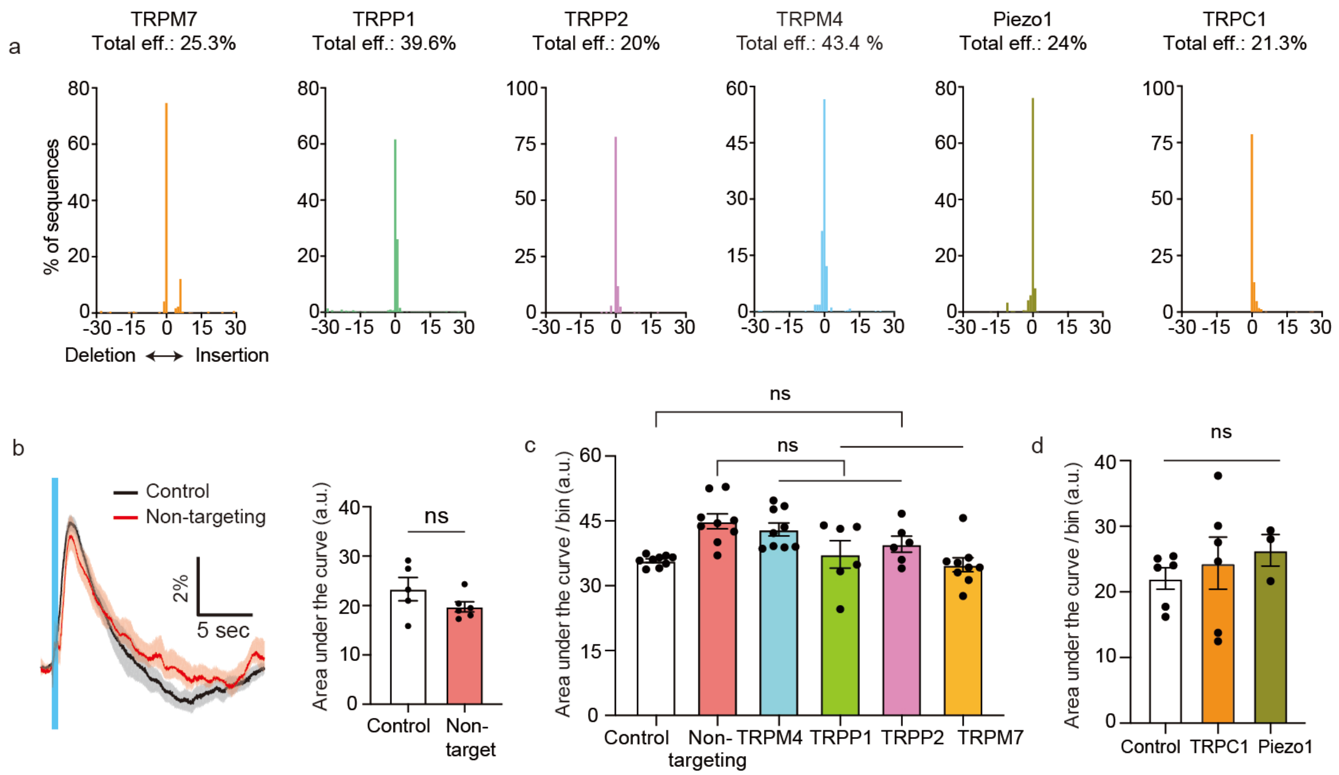


**Supplementary Figure 8 | CRISPR/Cas9 for knockdown specific mechanosensitive ion channel.** (a) CRISPR/Cas9 efficiencies for each targeted channel. (b) Calcium responses from wild type neurons and modified neurons (CRISPR/Cas9 with non-targeting sgRNA as a positive control (n= 5 dishes, Unpaired T-test, p=0.1720). (c and d) Quantification of spontaneous activities after knock down the target channels (n= 6-9 dishes each. P=0.0821~0.9999 (control vs channels), p=0.0024~0.9999 (Non-targeting vs channels), n= 6 dishes each, P=0.8051~0.9999 (control vs channels).

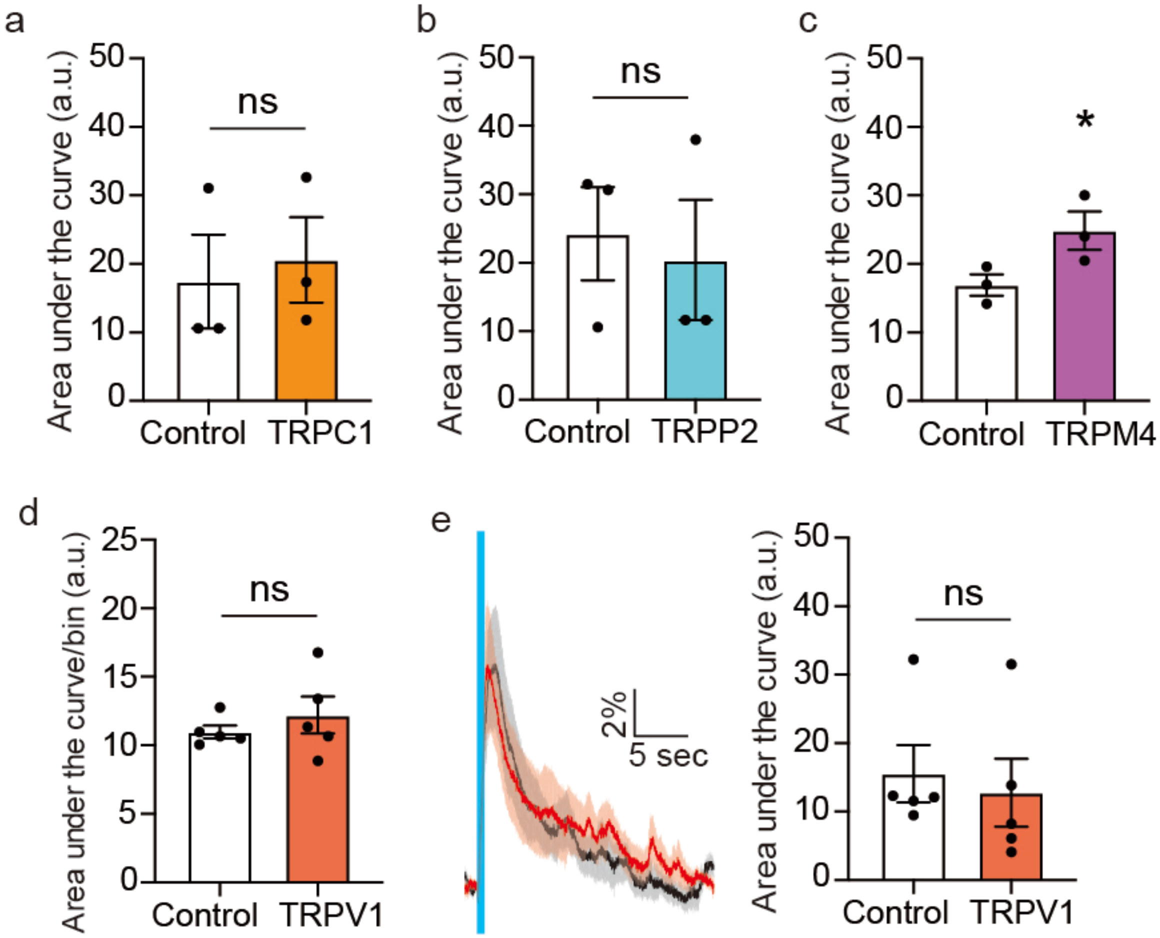


**Supplementary Figure 9 | Overexpression of mechanosensitive channels.** (a) Quantification of spontaneous activity after overexpressing TRPC1 (n= 3 dishes, Unpaired T-test, P=0.7491), (b) TRPP2 (n= 3 dishes, Unpaired T-test, P=0.7473), and (c) TRPM4 (n= 3 dishes, Unpaired T-test, P=0.0683). (d) Quantification of spontaneous activity after overexpressing TRPV1 (n= 5 dishes, Unpaired T-test, P=0.4191). (e) Calcium response after overexpressing TRPV1 (n= 5 dishes, Unpaired T-test, P=0.6801).

**Supplementary Movie 1 | Calcium responses of neurons to ultrasound stimulation.** Real time imaging of calcium responses from GCaMP6f neurons to ultrasound stimulation. Ultrasound intensity at 15 W/cm$^2$ was applied for 500 ms 2 times with 20 sec inter-pulse interval.

**Supplementary Table 1. sgRNAs for CRISPR knockdown of mechanosensitive channels**

| Target channel | sgRNA | PCR primers for seq. (For/Rev) |
|---|---|---|
| TRPM4 | ACACAGTGGCCGGATCCGTG | caatgtgtctccccattg/ gaaaccctgtcttaaaaaagc |
| TRPM7 | TCCAGGGTAATCTCCCCCCG | gatttgaactcaggacttcc/ gatggatctctatgttgcc |
| TRPP1 | GAACTTGGCATACGGCATCG | cattagctcaacacgctg/ gagtatacccttacaacccac |
| TRPP2 | CCAATGTGTACTACTACACT | ccaccttgctacacatttc/ cctgccattttagcactg |
| Piezo1 | TGGGGCTGGAGTAGTTAGGG | cttcagctaagcagatgag/ ctttccaataattccctgacc |
| TRPC1 | TCTTACAGGTGGGCTTACGG | cagagtggtagaacacttg/ ccacagtatgagaaggtaatc |
| Scramble | ACAACACGCCGACACGTCTA | |

**Supplementary Table 1. sgRNAs for CRISPR knockdown of mechanosensitive channels**

| Target channel | sgRNA | Off target DNA | Chrm.. | Position | Mismatching | CDF score |
|---|---|---|---|---|---|---|
| TRPM4 | ACACAGTGGCCGGATCCGTGNGG | | n/a | | | |
| TRPM7 | TCCAGGGTAATCTCCCCCCGNGG | | n/a | | | |
| TRPP1 | GAACTTGGCATACGGCATCGNGG | GAACTcGGCATAgGGCATCtAGG | chr7 | 58190249 | 3 | 0.086776859 |
| | | GAACgTGcCATACGGCATCcTGG | chrX | 7697030 | 3 | 0.168791209 |
| TRPP2 | CCAATGTGTACTACTACACTNGG | CaAATGaGTACcACTACACTGGG | chr8 | 86046724 | 3 | 0.602870813 |
| | | gCtATGTGTcCTACTACACTAGG | chr7 | 129894326 | 3 | 0.214833759 |
| | | CCAAaaTGTACTACaACACTGGG | chr4 | 107776116 | 3 | 0.289473684 |
| | | CCAATGTGTACTACcACcgTAGG | chr4 | 116068603 | 3 | 0.006493506 |
| | | CaAATGTGTACTACctCACTTGG | chr5 | 122374594 | 3 | 0.010017531 |
| | | CaAATGTaTACTACcACACTTGG | chr5 | 133111346 | 3 | 0.198347108 |
| | | gCAATGTGTACTACTACAagAGG | chr15 | 7216283 | 3 | 0.069053708 |
| | | CCAATGTGTAaTgCcACACTGGG | chr6 | 113238793 | 3 | 0.13339921 |
| | | CCAcTGTGTACcgCTACACTTGG | chr14 | 102951205 | 3 | 0.218064342 |
| | | CCAATGTGTACcACTACAtgGGG | chr9 | 110447439 | 3 | 0.077161229 |
| | | CCAtTGTGTAgTACTACAtTGGG | chrX | 61766440 | 3 | 0.07342657 |
| | | CCAcTtTGTACTACTACAtTTGG | chr11 | 25615191 | 3 | 0.108597285 |
| | | CaAATGTGTACcACcACACTGGG | chr11 | 65510726 | 3 | 0.187907786 |
| | | CaAATGTGTgCTACTACACTTGG | chr2 | 6718054 | 2 | 0.404040404 |
| **Piezo1** | TGGGGCTGGAGTAGTTAGGGNGG | TGGGGCaGGAGTAGgTgGGGTGG | chr8 | 38306807 | 3 | 0.007720588 |
| | | TGGGGCaGGAGTAGgTAGGtGGG | chr12 | 37700661 | 3 | 0.030625 |
| | | TGGGGCTGaAGTgGTaAGGGAGG | chr12 | 86241822 | 3 | 0.381140599 |
| | | TGGGGagGGAGTAGTgAGGGAGG | chr3 | 84110027 | 3 | 0.18907563 |
| | | TGGGGCTGGAGaAGcaAGGGTGG | chr7 | 109584451 | 3 | 0.198347108 |
| | | TGGGGCaGGAGTAGgTAGGtGGG | chr7 | 120173029 | 3 | 0.030625 |

| | | | | | | |
|---|---|---|---|---|---|---|
| | | TGGGGgTGGAGTAaTcAGGGAGG | chr7 | 139163261 | 3 | 0.25 |
| | | TGGGGCgGagGTAGTTAGGGTGG | chr1 | 39664346 | 3 | 0.210084034 |
| | | TGGGGaaGGAGTAGTTAGGaTGG | chr15 | 35582782 | 3 | 0.76171875 |
| | | TGaGGCTGGAGgAGaTAGGGAGG | chr15 | 101772220 | 3 | 0.217105263 |
| | | TGGGGCTGGAGcAGTgAGGcAGG | chr17 | 44247595 | 3 | 0.140543667 |
| | | TGaGGCTGGAGTAGgTgGGGGGG | chr10 | 26559790 | 3 | 0.006617647 |
| | | TGGGGCTGGAGaAGTgAGGcAGG | chr6 | 84393994 | 3 | 0.118681319 |
| | | TGGGGgTGGAGTtGgTAGGGTGG | chr6 | 98682092 | 3 | 0.0075 |
| | | TGGGGCTGcAcTAGTgAGGGAGG | chr6 | 110647213 | 3 | 0.079881657 |
| | | TGtGGCTGGAGTgGgTAGGGTGG | chr6 | 123922209 | 3 | 0.016304348 |
| | | TGGGGCTGGAGTctTTAGaGAGG | chr14 | 116081202 | 3 | 0.040100251 |
| | | TGGGGCTGGAGTgGgTAGGtGGG | chr9 | 36602467 | 3 | 0.022826087 |
| | | TGGGGgTGGAGaAGTTgGGGGGG | chr18 | 69502225 | 3 | 0.070588235 |
| | | TGGGGgTGGAGgAGTTgGGGTGG | chr11 | 116447021 | 3 | 0.044117647 |
| **TRPC1** | TCTTACAGGTGGGCTTACGGNGG | TCTTcCAGGTGGGaTTACGtTGG | chr12 | 33176485 | 3 | 0.1225 |
| | | TCTTACAGcTGatCTTACGGAGG | chr3 | 139139588 | 3 | 0.150769231 |
| | | TCTTACAGGTatGCTTACaGAGG | chr16 | 59945876 | 3 | 0.274725275 |
| | | TCTaACAcGTGaGCTTACGGGGG | chr1 | 189833763 | 3 | 0.273504273 |
